# Lupus autoantibodies initiate a maladaptive equilibrium sustained by HMGB1:RAGE signaling and reversed by LAIR-1:C1q signaling

**DOI:** 10.1101/2023.05.01.538935

**Authors:** Kaitlin R. Carroll, Mark Mizrachi, Sean Simmons, Bahtiyar Toz, Jeffrey Wingard, Nazila Tehrani, Aida Zarfeshani, Nina Kello, Czeslawa Kowal, Joshua Z. Levin, Bruce T. Volpe, Betty Diamond

## Abstract

Cognitive impairment is a frequent manifestation of neuropsychiatric systemic lupus erythematosus (NPSLE), present in up to 80% of patients and leading to a diminished quality of life. We have developed a model of lupus-like cognitive impairment which is initiated when anti-DNA, anti-N-methyl D-aspartate receptor (NMDAR) cross-reactive antibodies, which are present in 30% of SLE patients, penetrate the hippocampus. This leads to immediate, self-limited excitotoxic death of CA1 pyramidal neurons followed by a significant loss of dendritic arborization in the remaining CA1 neurons and impaired spatial memory. Both microglia and C1q are required for dendritic loss. Here we show that this pattern of hippocampal injury creates a maladaptive equilibrium that is sustained for at least one year. It requires HMGB1 secretion by neurons to bind RAGE, a receptor for HMGB1 expressed on microglia, and leads to decreased expression of microglial LAIR-1, an inhibitory receptor for C1q. The angiotensin converting enzyme (ACE) inhibitor captopril, which can restore a healthy equilibrium, microglial quiescence, and intact spatial memory, leads to upregulation of LAIR-1. This paradigm highlights HMGB1:RAGE and C1q:LAIR-1 interactions as pivotal pathways in the microglial–neuronal interplay that defines a physiologic versus a maladaptive equilibrium.

## Introduction

The autoimmune disease systemic lupus erythematosus (SLE) is characterized by the production of self-reactive antibodies leading to multi-organ system involvement and is frequently associated with neurological symptoms, most commonly cognitive dysfunction^1^. While the pathophysiology of NPSLE is not well understood, and indeed is likely caused by several distinct triggers and mechanisms, we have developed a model of cognitive dysfunction caused by anti-double-stranded DNA (dsDNA) antibodies which cross-react with the excitatory neuronal NMDAR^2,3^. These antibodies, termed DNRAbs, are present at high titers in approximately 30% of SLE patients and in a higher percentage of patients with NPSLE^4^. They have been found in brain tissue of SLE patients^5^, and their presence in cerebrospinal fluid (CSF) has been associated with non-focal NPSLE symptoms such as cognitive dysfunction^6^.

DNRAbs can be induced by immunizing mice with a peptide mimetope of DNA derived from a consensus sequence, DWEYS, which is shared by the GluN2A and GluN2B subunits of the NMDAR, and is multimerized on a polylysine backbone^2,7^. When lipopolysaccharide (LPS) is administered systemically several weeks post-immunization, there is a transient increase in permeability in the blood-brain barrier (BBB) allowing circulating immunoglobulin to penetrate the hippocampus^2^. DNRAbs, which act as positive allosteric modulators of the NMDAR^8,9^, cause excitotoxic death of approximately 30% of CA1 pyramidal neurons^2,9^. The remaining neurons that experience NMDAR activation that remains below the threshold for excitotoxicity secrete HMGB1^10,11^, a nuclear protein that, when secreted, can act as a damage-associated molecular pattern (DAMP)^12^ and can bind the NMDAR and potentiate NMDAR activation^10,13^ as well as directly impair neuronal function through other pathways^14,15^. Cytosolic HMGB1, a precursor to secreted HMGB1, is abundant in the hippocampus of mice immunized to produce DNRAbs followed by administration of systemic LPS (DNRAb^+^ mice) compared to control mice immunized with the polylysine backbone alone and given systemic LPS (DNRAb^−^ mice)^10^. There is no acute excitotoxicity and significantly less cytosolic HMGB1 in CA1 pyramidal neurons in DNRAb^−^ mice^10^.

Notably, we detect no microglial activation (based on morphology and CD68 expression^10,16^) in DNRAb^+^ mice two weeks following LPS administration, at which point DNRAbs are no longer detectable in the brain^10^. By eight weeks post-LPS, substantial microglial activation, a clear loss of dendritic arborization, and impaired spatial memory are observed in DNRAb^+^ mice. This pathology is mediated by microglia and the complement protein C1q and can be prevented by inhibitors of the angiotensin converting enzyme (ACE inhibitors)^10^. Importantly, this brain pathology is sustained for at least 12 months, the last time point at which DNRAb^+^ mice have been assessed, demonstrating a persistent state of neuroinflammation long past the transient exposure to DNRAbs and the acute damage they induce (Supp Fig 1). Here we investigate the mechanisms of this sustained neuroinflammation and the restoration of a healthy homeostasis by ACE inhibitors.

## Results

### Neuronal HMGB1 induces microglial activation through RAGE

The chromatin-binding protein HMGB1 can be secreted by activated, stressed, or damaged cells to act as a DAMP and carry nucleic acid ligands to endosomal toll-like receptors (TLRs) to activate monocytes and macrophages^17^. We asked whether HMGB1 plays a role in inducing the reactive microglial state characteristic of DNRAb^+^ mice, given the substantial increase in cytosolic HMGB1 in these mice^10^. We generated neuronal HMGB1-deficient (Camk2a-cre^+^ HMGB1^fl/fl^, termed HMGB1 cKO) DNRAb^+^ mice and observed significantly decreased microglial activation in HMGB1 cKO mice compared with their HMGB1-sufficient counterparts (Camk2a-cre^-^ HMGB1^fl/fl^; Fig 1A-B). To determine the direct effect of HMGB1 on microglia, we cultured primary microglia *in vitro* with HMGB1 and observed a dose-dependent increase in mRNA expression of proinflammatory cytokines (*Tnf, Il1b*) and *C1qa* (Fig 1C), as well as increased secretion of TNFα and IL-1β (Fig 1D). HMGB1 also induced type I interferon (IFN; *Ifnb*) expression as well as IFNβ secretion and upregulation of genes in the IFN pathway such as *Irf7* and the IFN-stimulated gene *Mx1* (Fig 1E-F). Notably we also detected increased IFNβ in the hippocampus of DNRAb^+^ compared to DNRAb^-^ mice (unpublished data). To further address the stimulus for increased expression of C1q, we cultured microglia with IFNβ and observed a significant increase in *C1qa* expression (Fig 1G). Together, these data indicate that neuronal HMGB1 induces microglial activation and secretion of proinflammatory cytokines and type I IFN, which, in turn, function to enhance C1q expression.

**Figure 1.**
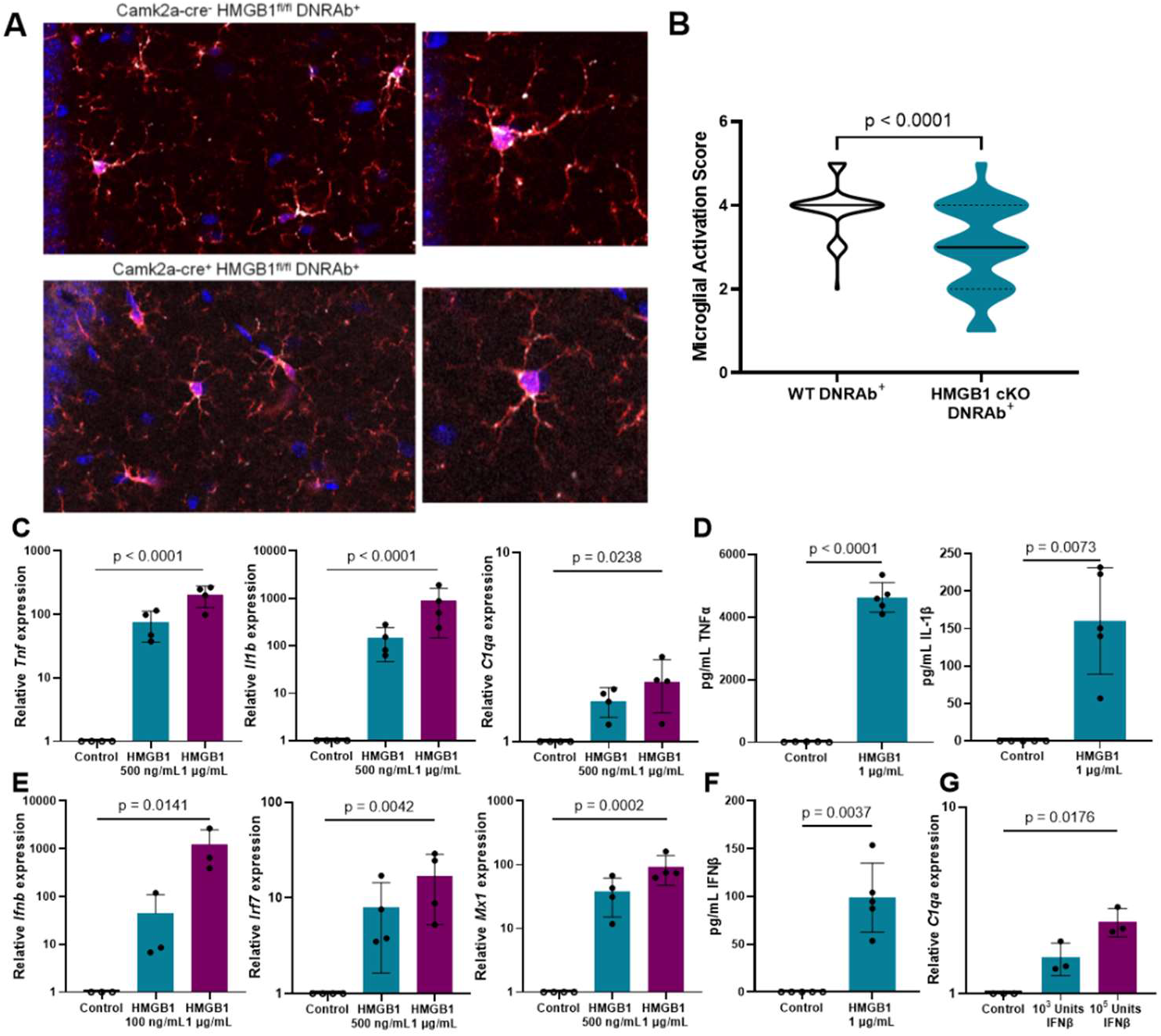
Microglia are activated by neuronal HMGB1. **A)** Representative sections of microglia in CA1 stratum radiatum stained for Iba1 (red) and CD68 (white) in Camk2a-cre^-^ HMGB1^fl/fl^ B6.H2^d^ DNRAb^+^ mice (WT; n=5, top) and Camk2a-cre^+^ HMGB1^fl/fl^ B6.H2^d^ DNRAb^+^ mice (HMGB1 cKO; n=5, bottom). **B)** Decreased activation score in microglia from Camk2a-cre^+^ HMGB1^fl/fl^ B6.H2^d^ (HMGB1 cKO) DNRAb^+^ mice compared to Camk2a-cre^-^ HMGB1^fl/fl^ B6.H2^d^ (WT) DNRAb^+^ mice based on morphology and CD68 expression (median (solid line) with quartiles (dash); n=5 mice per group; n=65 microglia scored per group; Mann-Whitney test). **C)** Relative mRNA expression of *Tnf, Il1b*, and *C1qa* is increased with HMGB1 stimulation in cultured WT (B6) microglia stimulated with 0, 500, or 1000 ng/ml HMGB1 for 4.5 hours in serum-free media (mean +/-SD; microglia cultured from 4 independent litters; one-way repeated measures ANOVA on log transformed data). **D)** Secretion of TNFα and IL-1β is increased with HMGB1 stimulation in cultured WT (B6) microglia stimulated with and without 1 μg/ml HMGB1 for 24 hours in serum-free media (mean +/-SD; microglia cultured from 5 independent litters; paired t-test). **E)** Relative mRNA expression of *Ifnb, Irf7*, and *Mx1* is increased with HMGB1 stimulation in cultured WT (B6) microglia stimulated with 0, 100, 500, or 1000 ng/ml HMGB1 for 4.5 hours in serum-free media (mean +/-SD; microglia cultured from 3-4 independent litters; one-way repeated measures ANOVA on log transformed data). **F)** Secretion of IFNβ is increased with HMGB1 stimulation in cultured WT (B6) microglia stimulated with and without 1 μg/ml HMGB1 for 24 hours in serum-free media (mean +/-SD; microglia cultured from 5 independent litters; paired t-test). **G)** Relative mRNA expression of *C1qa* is increased with IFNβ stimulation in cultured WT (B6) microglia stimulated with 0, 10^3^, or 10^5^ Units IFNβ for 4.5 hours in serum-free media (mean +/-SD; microglia cultured from 3 independent litters; one-way repeated measures ANOVA on log transformed data).

HMGB1 binds to numerous cell surface receptors, including the receptor for advanced glycation end-products (RAGE)^17^. Crucially, when HMGB1 binds to RAGE, this complex chaperones nucleic acids to endosomal TLRs and leads to upregulation of both inflammatory cytokines and type I IFN^18^. Therefore, we asked whether RAGE expression is necessary for microglial activation in this model of NPSLE (Fig 2). We incubated microglia from RAGE– deficient (RAGE KO) mice with HMGB1 and found that the transcription of *Tnf, Il1b, Ifnb, Irf7*, and *Mx1* was significantly attenuated in RAGE KO microglia compared with WT (Fig. 2A). We then asked whether microglial activation would occur *in vivo* in RAGE KO DNRAb^+^ mice. DNRAbs are induced by peptide immunization only in H-2^d^ mice^19^. As we were unable to generate a C57BL/6 mouse with a recombination event that rendered it RAGE-deficient on an H-2^d^ background due to proximity of the genes encoding RAGE and MHC, we injected C57BL/6 WT mice and C57BL/6 RAGE KO mice with G11, a monoclonal DNRAb (mDNRAb^+^), or B1, an isotype matched monoclonal antibody with no detectable binding in the brain (mDNRAb^−^)^20^, followed by LPS administration to mediate hippocampal BBB breach. After 8 weeks, the CA1 region of the hippocampus was examined in each strain. While acute pyramidal neuron loss occurred in both RAGE KO DNRAb^+^ mice and WT DNRAb^+^ mice (Supp Fig 2A), RAGE KO mice developed neither microglial activation (Fig 2B-C) nor decreased dendritic complexity, while this pathology was observed in WT DNRAb^+^ mice (Fig 2D-E), demonstrating that RAGE expression is necessary for the development of the sustained state of neuroinflammation caused by DNRAbs.

**Figure 2.**
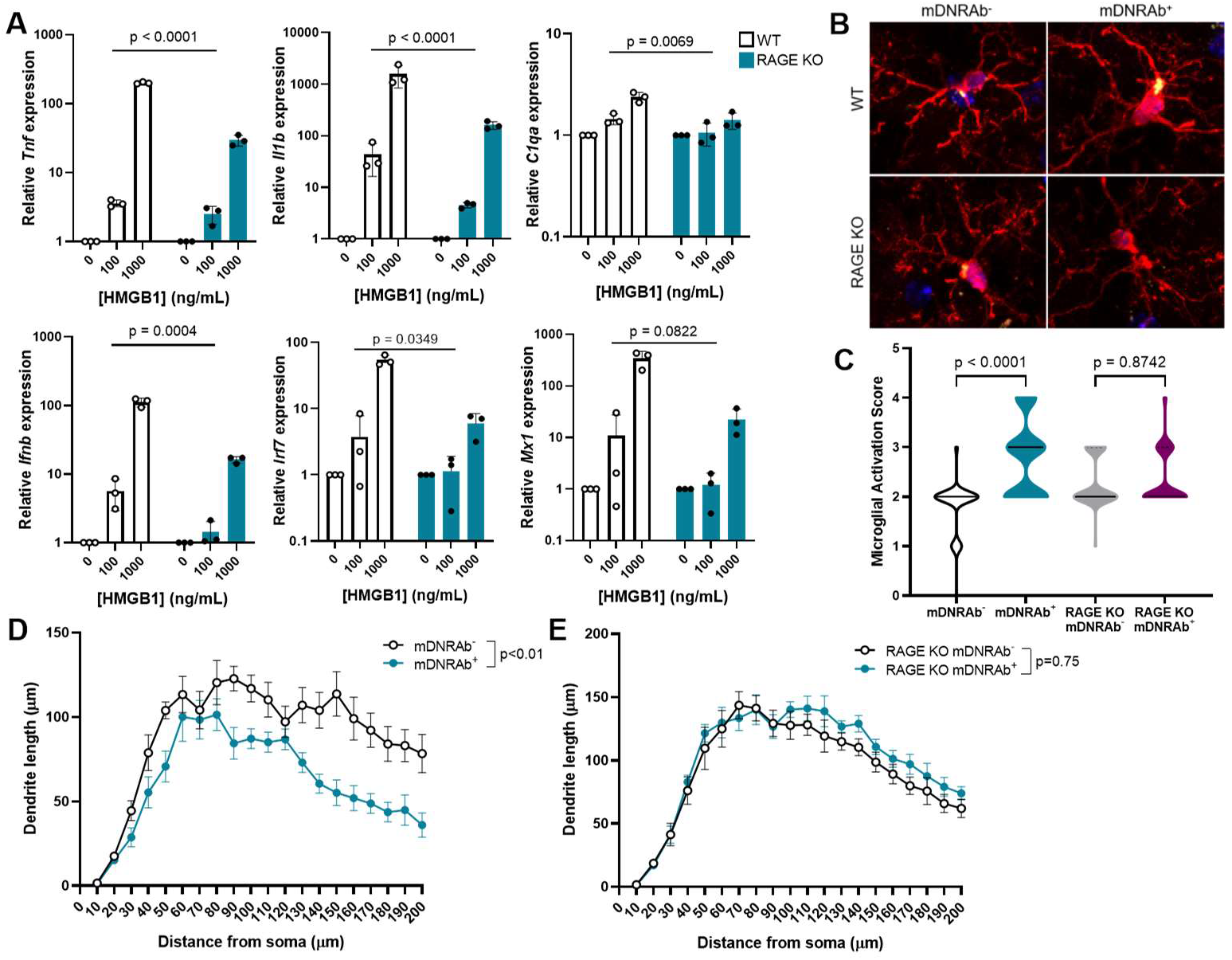
DNRAbs induce microglial activation through RAGE. **A)** Loss of RAGE decreases the relative mRNA expression of *Tnf, Il1b, C1qa, Ifnb, Irf7*, and *Mx1* with HMGB1 stimulation compared with WT (B6) microglia in WT and RAGE KO cultured microglia stimulated with 0, 100, or 1000 ng/ml HMGB1 for 4.5 hours in serum-free media (mean +/-SD; microglia cultured from 3 independent litters; two-way repeated measures ANOVA on log transformed data). **B)** Representative sections of microglia in CA1 stratum radiatum stained for Iba1 (red) and CD68 (yellow) in WT (B6) and RAGE KO mDNRAb^-^ (left) and mDNRAb^+^ (right) mice (n=5 mice per group). **C)** Increased activation score in microglia in DNRAb^+^ WT (B6) but not RAGE KO mice compared to DNRAb^-^ counterparts based on morphology and CD68 expression (median (solid line) with quartiles (dash); n=5 mice per group; n=74-87 microglia scored per group; Kruskal-Wallis test). **D)** Analysis of dendritic complexity shows a decrease in WT (B6) mDNRAb^+^ compared to mDNRAb^-^ groups (mean +/-SEM; n=7 mice per group; n=25 neurons analyzed per group; linear mixed model test). **E)** Analysis of dendritic complexity shows no difference in RAGE KO mDNRAb^+^ compared to mDNRAb^-^ groups (mean +/-SEM; n=6-7 mice per group; n=24 neurons analyzed per group; linear mixed model test).

### Treatment effects of ACE inhibitors depend on LAIR-1 expression

We have shown that in the periphery C1q can act as a modulating agent tempering the proinflammatory effects of HMGB1:RAGE signaling in monocytes^18^. While complement frequently plays an anti-inflammatory role outside the brain, its function in the central nervous system (CNS) is more complex and highly contextual^10,16,21,22^. C1q plays a pivotal role in synaptic pruning during development by tagging synapses for elimination^21^. Moreover, C1q appears to continue to play a role in synaptic remodeling throughout adulthood, and increasing evidence reveals that this process becomes maladaptive in the context of aging and brain injury^22,23^. For instance, Alzheimer’s disease and other neurodegenerative diseases show increases in complement proteins in the brain and CSF^22^. Although C1q appears to contribute to a sustained maladaptive equilibrium in the CNS and is critical in the DNRAb-induced model of NPSLE, we reasoned that C1q might also mitigate microglial activation through binding the inhibitory receptor LAIR-1, as occurs in peripheral macrophages^18^.

We further hypothesized that LAIR-1 might be reduced on hippocampal microglia of DNRAb^+^ mice and increased following treatment with an ACE inhibitor. We therefore assessed the expression of LAIR-1 in hippocampal microglia by qRT-PCR and found that microglia from DNRAb^+^ mice indeed expressed significantly less *Lair1* transcript than microglia from DNRAb^−^ mice (Fig 3A). In addition, we treated DNRAb^+^ mice with ACE inhibitors, captopril which penetrates the BBB, or enalapril which does not^10^. We observed increased *Lair1* expression in hippocampal microglia isolated from captopril-treated mice compared to enalapril-treated mice, suggesting that LAIR-1 might indeed be an important regulator of microglial activation state (Fig 3B).

**Figure 3.**
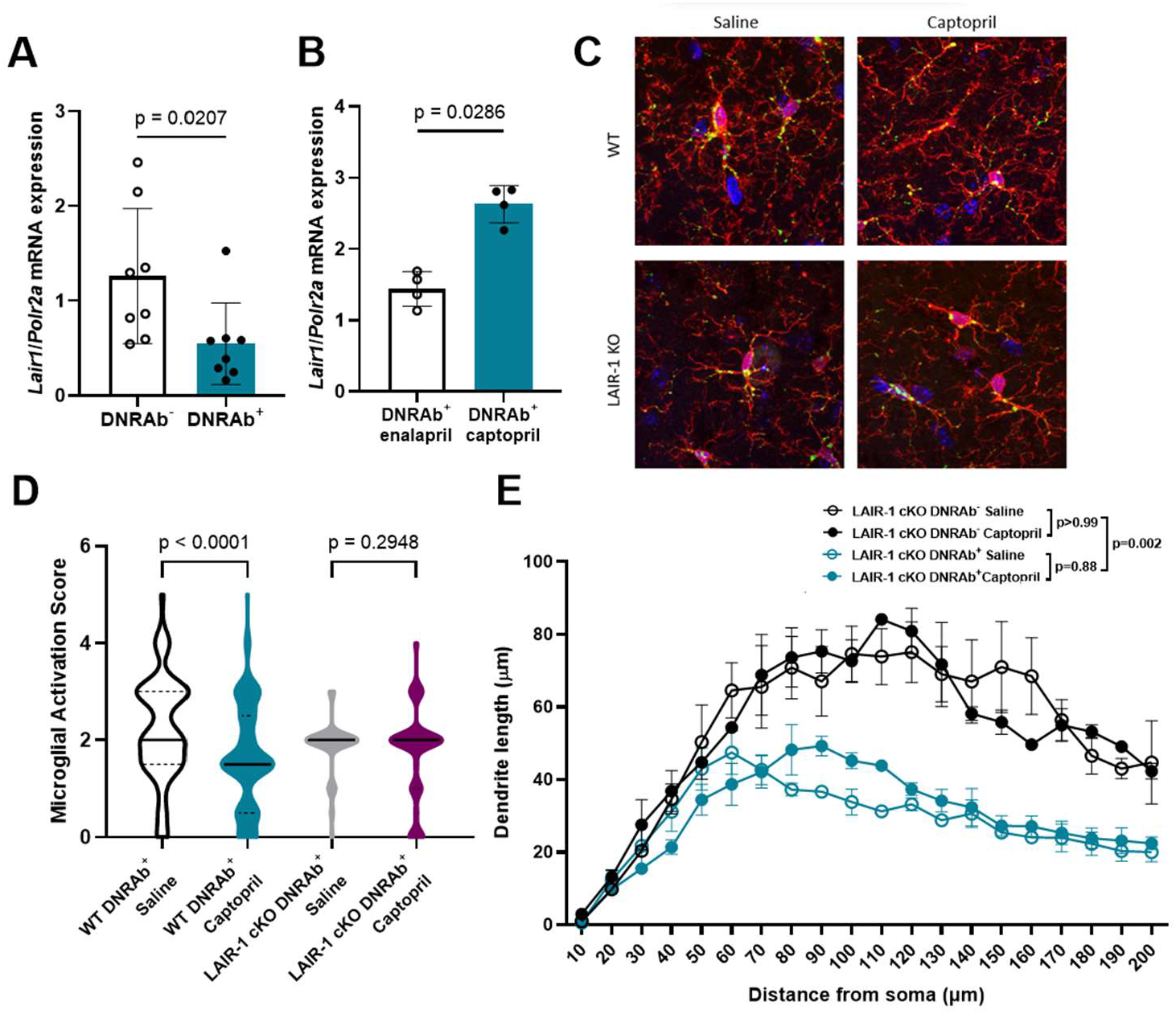
Treatment with ACE inhibitors requires LAIR-1 for efficacy. **A)** Decreased *Lair1* mRNA expression in microglia isolated from DNRAb^+^ compared with DNRAb^-^ B6.H2^d^ mice (mean +/-SD; n=8 mice per group, pooled from two independent experiments; Mann-Whitney test). **B)** *Lair1* mRNA expression in microglia isolated from DNRAb^+^ B6.H2^d^ mice is decreased in mice treated with captopril (5 mg/kg i.p.) compared with those treated with enalapril (5 mg/kg i.p.) daily for 2 weeks (mean +/-SD; n=4 mice per group; Mann-Whitney test). **C)** Representative sections of microglia in CA1 stratum radiatum stained for Iba1 (red) and CD68 (green) in DNRAb^+^ WT (B6.H2^d^) and LAIR-1 cKO mDNRAb^-^ mice treated with saline (left) or captopril (5 mg/kg, right; n=5 mice per group). **D)** Decreased activation score in DNRAb^+^ microglia in WT (B6.H2^d^) but not LAIR-1 cKO mice treated with captopril compared to saline-treated counterparts based on morphology and CD68 expression (median (solid line) with quartiles (dash); n=3-5 mice per group; n=70-180 microglia scored per group; Kruskal-Wallis test). **E)** Captopril treatment has no effect on dendritic complexity in LAIR-1 cKO DNRAb^-^ and DNRAb^+^ mice treated with either saline or captopril (5 mg/kg; mean +/-SEM; n=2-3 mice per group; n=19-27 neurons analyzed per group; linear mixed model test).

To test whether the decrease in microglial activation and the increased neuronal dendrite arborization mediated by captopril are dependent on LAIR-1, we immunized WT mice and mice with LAIR-1 deficiency in microglia (Lyz2-cre^+^ LAIR-1^fl/fl^; termed LAIR-1 cKO). Both strains of DNRAb^+^ mice developed acute neuronal loss in the hippocampal CA1 pyramidal layer (Supp Fig 2B) and exhibited increased microglial activation and loss of dendritic arborization (Fig 3C-E). While captopril treatment led to a lower activation score for microglia in WT mice, it did not affect the microglial activation state in LAIR-1 cKO mice (Fig 3C-D).

Moreover, while captopril ameliorated the loss of dendritic complexity in WT DNRAb^+^ mice, it had no effect on dendritic arborization in LAIR-1 Cko DNRAb^+^ mice (Fig 3E). Together, these observations demonstrate that microglial LAIR-1 is essential for the regulation of inflammation and neuroprotective effects of ACE inhibitors following DNRAb-mediated neuronal injury.

### Transcriptional profiling of hippocampal cells in DNRAb^+^ and DNRAb^-^ mice

To further interrogate the pathways activated by DNRAb in the hippocampus, we performed single-cell (sc) RNA-seq on hippocampal microglia from saline-treated DNRAb^-^, saline-treated DNRAb^+^, and captopril-treated DNRAb^+^ mice. We identified six clusters of microglia by scRNA-seq (Fig 4A), including two large clusters, one expressing genes characteristic of a healthy, quiescent homeostatic state (Homeostatic) and one with high levels of the transmembrane chemosensor *Ms4a7* (Ms4a7+)^24^. We also identified four very small clusters: one with *Top2a*+ cycling cells (Cycling), one with *S100a4*+ microglia (S100a4+), one with high expression of IFN-stimulated genes (IFN-responsive)^24^, and one with very low expression of *Tmem119* (Tmem119-) which may be phagocytic microglia or macrophages^25-28^.

**Figure 4.**
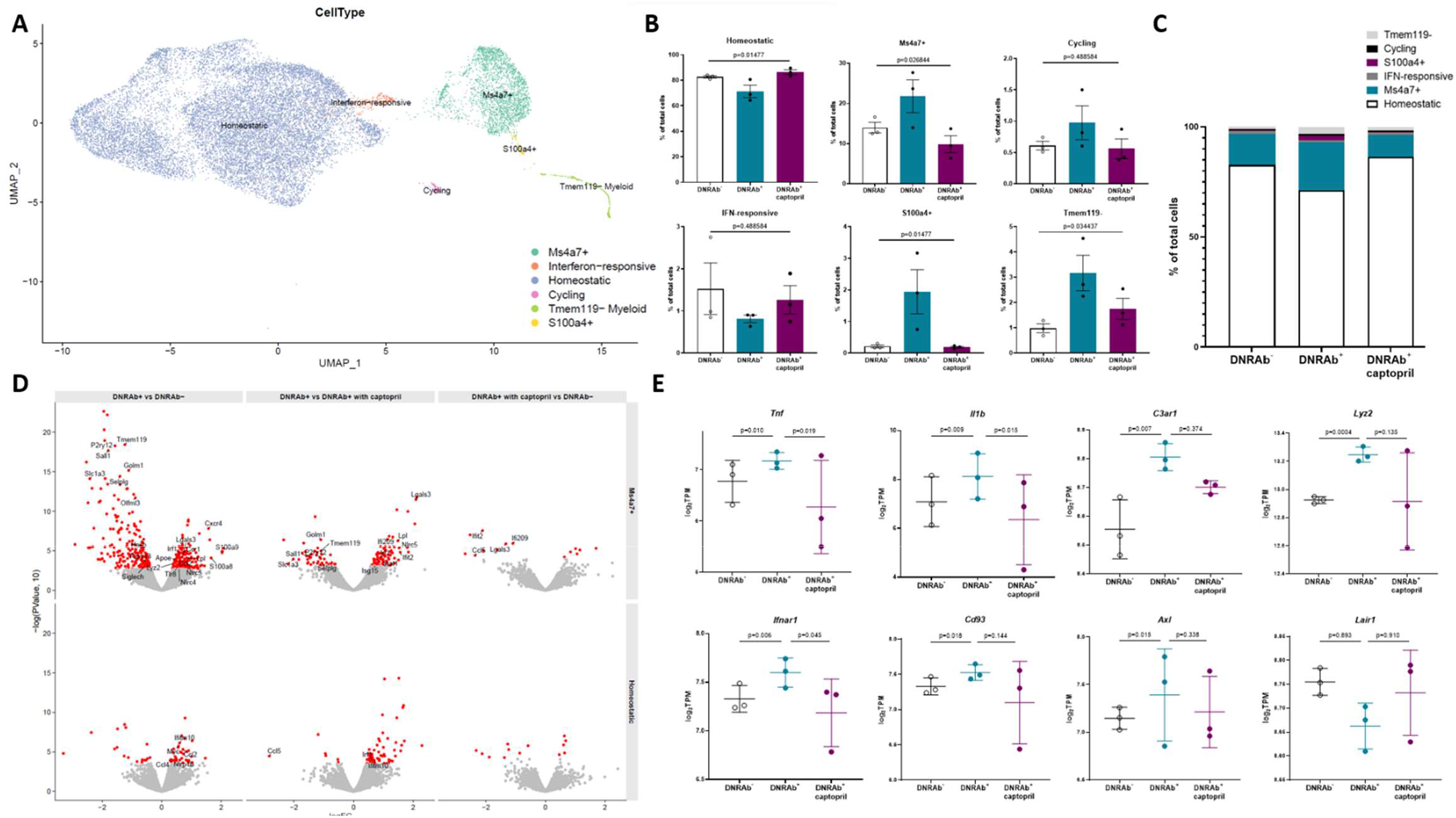
Microglial clustering in single-cell RNA-seq shows captopril treatment ameliorates DNRAb-induced inflammatory phenotype. **A)** UMAP plot of single-cell microglia data, with each cell colored by microglia cluster: Cycling (pink), Homeostatic (blue), IFN-responsive (orange), Ms4a7+ (green), S100a4+ (yellow), Tmem119-(light green). **B)** Cell type composition plots; percent of total cells represented in each cluster in DNRAb^-^, DNRAb^+^, and captopril-treated DNRAb^+^ mice (mean +/-SEM; n=3 mice per group; ANOVA FDR adjusted p-values shown). **C)** Stacked bar plots of the cell type composition in each condition. **D)** Volcano plots of the DE results comparing between different conditions (columns) in Ms4a7+ and Homeostatic microglia (rows). The y-axis is log base 10 p-value, the x axis is the log fold change (logFC) reported by EdgeR. Genes with FDR < 0.05 are colored red, and genes of interest are labeled. **E)** Expression of genes of interest in different conditions in Ms4a7+ microglia. Pseudobulk expression levels are calculated on a per mouse basis (y axis, log base 2 transcripts per million), stratified by condition (mean +/-SD; p-values are uncorrected).

We focused primarily on the two larger clusters of microglia, the Homeostatic and Ms4a7+ clusters, as several mice provided fewer than 20 cells to each of the other clusters. Our analysis revealed significant differences in the proportion of each cluster between treatment groups (Homeostatic p=0.004, adjusted p=0.01; Ms4a7+ p=0.013, adjusted p=0.0268; one-way ANOVA) demonstrating that DNRAb^+^ mice had an increased proportion of activated microglia and that captopril treatment returns the proportions to control levels (Fig 4B-C, Supp Tables 2-3). Interestingly, the transcriptional profile of Ms4a7+ microglia displays similarity to those of microglia with dysregulated functionality from multiple models of neuroinflammation and neurodegeneration, including the disease-associated microglia (DAM)^26^, NPSLE^29^, and neurodegenerative microglia (MGnD)^30^ gene signatures (Supp Fig 4).

Comparison of differentially expressed genes (DEGs) among the experimental groups demonstrated that Homeostatic microglia were qualitatively comparable in all groups, but Ms4a7+ microglia were increased both in their relative quantity and in gene expression differences in DNRAb^+^ mice (Fig4B-D, Supp Tables 2-5). Ms4a7+ microglia from DNRAb+ mice exhibited upregulation of genes associated with inflammation, phagocytosis, and the DAM signature (*Cxcr4, Lgals3, Apoe, Ccl2, Lyz2, Tlr8, Nlrc5*) and downregulation of genes indicative of a more quiescent, basal state (*P2ry12, P2ry13, Sall1, Selplg, Olfml3, Siglech, Hexb*) compared to Ms4a7+ microglia from DNRAb^-^ and captopril-treated DNRAb^+^ mice (Fig 4D)^26^. In stark contrast, few genes were differentially expressed between DNRAb^-^ mice and captopril-treated DNRAb^+^ mice in either cluster (Fig 4D). Based on our *in vitro* and histological data (Figs 1-3), we analyzed Ms4a7+ microglia from DNRAb^+^ and DNRAb^-^ mice for differential expression in inflammatory cytokines, type I IFN, complement, and phagocytic pathways. Consistent with these data, Ms4a7+ microglia from DNRAb^+^ mice had increased expression of the proinflammatory cytokines *Tnfa* and *Il1b*; the type I IFN receptor *Ifnar1*; receptors for complement components such as *C3ar1* and *Cd93*; and the phagocytosis-associated receptor tyrosine kinase *Axl* and lysosomal protein *Lyz2* compared with DNRAb^-^ microglia (Fig 4E, Supp Tables 4-5). These changes were either attenuated or abrogated completely in captopril-treated DNRAb^+^ mice (Fig 4E, Supp Tables 4-5).

## Discussion

We have demonstrated that neuronal secretion of HMGB1 following DNRAb-mediated neuronal injury activates microglia and causes them to secrete proinflammatory cytokines and type I IFN, a process that depends upon RAGE expression in microglia. HMGB1 acts not only as a DAMP to activate microglia but also binds the NMDAR on neurons and can potentiate NMDAR signaling^13^. The secreted HMGB1 and type I IFN both induce microglia to upregulate expression of the complement component C1q, which binds neuronally derived HMGB1, thereby decorating HMGB1-bound synaptic proteins^10^. This complex likely acts as an opsonin on neuronal synapses to induce aberrant dendritic pruning and ultimately a loss of dendritic arborization and impaired spatial memory characteristic of this pathology^10^. Following ACE inhibition, microglia return to a quiescent state and deleterious dendritic loss ceases, a process that depends upon microglial expression of the inhibitory receptor LAIR-1.

While LAIR-1 increases with ACE inhibition and is required for its efficacy in this context, it will be important to determine the mechanisms underlying this effect. ACE inhibitors affect both the renin-angiotensin and kallikrein-kinin systems^31,32^. ACE converts angiotensin I to angiotensin II, which can induce a proinflammatory phenotype in microglia when it binds to angiotensin receptor 1^32^. ACE also degrades bradykinin, which would otherwise exert anti-inflammatory effects on microglia^33^. Moreover, bradykinin has been reported to decrease type I IFN production in bone marrow-derived dendritic cells^31,32^ and captopril has been shown to decrease type I IFN production and improve behavioral symptoms in other mouse models of NPSLE^34^. It will thus also be crucial to determine whether IFN is necessary for pathogenesis in this model and to ask whether the effects of captopril depend also upon a decrease in IFN and pro-inflammatory cytokines.

We have also shown through transcriptomic analysis that hippocampal microglia can largely be separated into two distinct clusters in this model; the Homeostatic cluster expresses genes characteristic of quiescent homeostatic state and the Ms4a7+ cluster expresses genes associated with a more reactive microglial phenotype and which resemble microglial gene signatures described in other neuroinflammatory contexts, including DAM^26^, NPSLE^29^, and MGnD^30^ microglia. These transcriptomic similarities may provide insight into shared pathogenic mechanisms. Additionally, while the Homeostatic microglia are proportionately decreased in DNRAb^+^ mice, they appear to be transcriptionally similar in all groups. Microglia from the Ms4a7+ cluster are not only proportionately increased in DNRAb^+^ mice, they are transcriptionally distinct from DNRAb^-^ and captopril-treated DNRAb^+^ Ms4a7+ microglia and reflect a more reactive and inflammatory state that is consistent with the maladaptive equilibrium that we observe in this phenotype.

Although the precipitating event of DNRAb penetration into the brain and the neuronal death it causes is transient, the ensuing state of microglial activation and loss of dendritic complexity is persistent. We hypothesize that the diminished number of synapses in the surviving neurons may increase NMDAR signaling strength such that there is greater synaptic activity in the damaged neurons leading to more secretion of HMGB1, initiating a self-reinforcing cycle as HMGB1 potentiates NMDAR signaling^10,13^. Interestingly, increased NMDAR signaling in neurons with fewer synapses has been described in a mouse model of schizophrenia^35^. In our model, increased extracellular HMGB1 induces the secretion of proinflammatory cytokines, type I IFN, and C1q. The inflammatory environment and loss of synapses, likely through complement-dependent microglial pruning, creates a self-perpetuating state of increased synaptic activity and HMGB1 secretion. Thus, neuronal stress, HMGB1 secretion, microglial activation, and loss of dendrites becomes a new, albeit maladaptive, equilibrium. This cycle can be interrupted, however, by the ACE inhibitor captopril as it enables microglia to express sufficient LAIR-1. Under this condition, microglia can return to a state of quiescence such that a healthy homeostasis can be regained, leading to restored dendritic and spine complexity and, remarkably, to amelioration of impaired spatial cognition^10^. Thus, HMGB1:RAGE and C1q:LAIR interactions determine the health of the brain after a lupus-like insult.

This model informs our understanding of NPSLE. Subjects with cognitive dysfunction exhibit specific alterations in the hippocampus, such as hypermetabolism indicative of increased synaptic activity, microglial activation, and BBB impairment, which are all reproduced faithfully in the murine model^36^; the data presented here suggest that neuroinflammation may be continue long after the triggering insult and that patients with quiescent systemic disease may still exhibit neuroinflammation. Altering RAGE activation and LAIR-1 expression using ACE inhibition may be of therapeutic benefit in patients with NPSLE, and perhaps in other individuals with ongoing, sustained neuroinflammation.

## Methods

### Animals

Mice were housed in the Center for Comparative Physiology at the Feinstein Institutes for Medical Research. C57BL/6 (B6) and B6.C-H2d/bByJ (B6.H2^d^) mice were purchased from The Jackson Laboratory. RAGE KO mice were a generous gift from Kevin Tracey, MD, of the Feinstein Institutes for Medical Research. LAIR-1 cKO mice were generated by crossing B6.129P2-Lyz2^tm1(cre)^Ifo/J with B6.Cg-Lair1^tm1Jco^/J (The Jackson Laboratory) after backcrossing onto the B6.H2^d^ background. B6.Cg-Tg(Camk2a-cre)T29-1Stl/J and Hmgb1^tm1.1Rshw^/J (Jackson Laboratories) were crossed with in house B6.H2^d^ mice to generate Camk2a-cre HMGB1^fl/fl^ B6.H2^d^ mice. All procedures using mice were conducted using protocols approved by The Feinstein Institutes of Medical Research IACUC (Protocols 2009-048 and 2022-023).

### Treatment

Female mice were immunized as previously described^2,10^. Briefly, mice were injected i.p. beginning at 6-8 weeks old with DWEYS peptide multimerized on a polylysine backbone (DNRAb^+^) or with the polylysine backbone alone (DNRAb^-^) in a Complete Freund’s Adjuvant (CFA; 263810, BD Biosciences) emulsion and boosted with two Incomplete Freund’s Adjuvant (IFA; 263910, BD Biosciences) emulsions spaced two weeks apart. Two weeks after the last immunization, mice were injected i.p. with 5 mg/kg body weight LPS (Sigma-Aldrich L4524, Lot#107M4048V) 48 hours apart. B6 and RAGE KO mice were passively immunized with 400 μg injected i.v. of the monoclonal antibody G11 (mDNRAb^+^) or B1 (mDNRAb^-^), produced as described previously^20^. Mice treated with an ACE inhibitor, either captopril (C8856, Sigma Aldrich) or enalapril (1235300, USP), once daily i.p. at 5 mg/kg body weight for two weeks beginning five weeks after the second LPS injection. Control mice received an equivalent volume of sterile saline based on body weight.

### Isolation of hippocampal microglia

Mice were anesthetized with Euthasol (Virbac) and perfused with pre-perfusion buffer containing 0.5% sodium nitrite, 0.9% sodium chloride, and 0.1% heparin. The hippocampus was extracted and dissociated using the Neural Tissue Dissociation Kit (Miltenyi Biotec), according to manufacturer’s instructions. Microglia were isolated according to manufacturer’s instructions using Myelin Removal Beads II (Miltenyi Biotec). Microglia used for qRT-PCR analysis were further sorted using CD11b MicroBeads (Miltenyi Biotec) and immediately lysed for RNA extraction. Microglia used for single-cell RNA-seq analysis were FACS sorted using Live/Dead Fixable Violet (Life Technologies L34964), FITC rat anti-mouse CD45 (Clone 30-F11, BD Biosciences), and PE rat anti-mouse CD11b (Clone M1/70, BD Biosciences).

### Primary microglia culture preparation

Primary microglia were isolated from P0-3 C57BL/6 or RAGE KO pups using the shaking method, adapted from previously described protocols^37^. Briefly, P0 litters were euthanized via decapitation, and brains were removed and collected under aseptic conditions. Brain tissue was dissociated to a single cell suspension using the neural tissue dissociation kit (Miltenyi Biotec), and the resulting cell suspension was centrifuged at 400g for 10 minutes. The cells were then resuspended in complete DMEM (with 4.5 g/L glucose) containing 10% FBS, 1% penicillin-streptomycin, and 0.5 ng/ml recombinant mouse GM-CSF (R&D Biosystems, Minneapolis, USA) before being filtered through a 70 μm cell strainer (431751, Corning). Cell suspensions from each litter of pups were plated into a different 175 cm^2^ flask and cultured until confluence (7-10 days). Microglia were recovered from culture by manual shaking. Growth medium containing ‘shaken’ microglia was centrifuged at 400g for 10 minutes, then counted and plated alone in a 48-well plate at 200,000 cells per well. Flow cytometry analysis using antibodies against CD45 (Clone 30-F11, BioLegend), CD11b (Clone M1-70, BD Biosciences), and TMEM119 (Clone 106-6, Abcam) indicated a microglial purity of greater than 95%.

### Cell culture treatment

Primary microglia were allowed to attach overnight in serum-free X-VIVO medium (Lonza Bioscience) before being treated with HMGB1 (100, 500, or 1000 ng/ml) or IFNβ (1000 or 100,000 Units). HMGB1 was obtained as a generous gift from Kevin Tracey, MD, of the Feinstein Institutes for Medical Research. IFNβ was purchased from R&D Biosystems (8234-MB-010/CF). Cells were left to incubate in treatments for 4.5 hours for analysis of lysate by qRT-PCR, and 24 hours for analysis of cell culture supernatant by ELISA.

### Quantitative Real-time PCR

Total RNA was extracted from microglia using the Qiagen RNeasy RNA extraction kit (Qiagen) according to the manufacturer’s instructions. Briefly, cells from each well were homogenized in RLT lysis buffer. The homogenate was then passed through QIAshredder spin columns to remove cellular debris, and RNA was purified using RNeasy spin columns. RNA was eluted in 30 μl ddH_2_0, and the quality and quantity of RNA were measured using NanoDrop. Reverse transcription of RNA was performed using the iScript Reverse Transcription Mix (Bio-Rad) according to the manufacturer’s protocol. Briefly, 15 μL of RNA was mixed with a reaction mixture including iScript Reverse Transcriptase enzyme, and the reaction was incubated at the recommended temperature cycle for cDNA synthesis. Quantitative real-time PCR (qRT-PCR) was performed using TaqMan Gene Expression Assays (Thermo Fisher Scientific) on a Roche LightCycler 480. Reactions were performed according to the manufacturer’s instructions. Gene expression levels were normalized to the expression of *Polr2a* using the ΔCt method. Relative expression levels were calculated using the ΔΔCt method.

### ELISA

Cell culture supernatant was collected from 24-hour treated microglia and spun down at 500g for 5 minutes followed by separation of supernatant. The Duoset TNFα, Duoset IL-1β, and Duoset IFNβ ELISAs (R&D Biosystems) were performed on supernatant according to the manufacturer’s instructions. Briefly, 96-well microplates were coated with the appropriate capture antibody and incubated overnight at 4° C. Plates were washed and blocked with 1% BSA for 1 hour. Samples and standards were added to the wells and incubated for 2 hours. After washing, detection antibody was added and incubated for 2 hours. Streptavidin-HRP conjugate was added and incubated for 20 minutes. The wells were washed, and substrate solution was added for 30-minute incubation. Stop solution was added to terminate the reaction, and the absorbance was read at OD450 using a microplate reader. Data was analyzed using a standard curve generated from the known standards provided in the kit, and statistical analysis was performed using GraphPad Prism.

### Immunohistochemistry

Mice were anesthetized with Euthasol (Virbac) or isofluorane (1.5%; Henry Schein Animal Health NDC11695-6776-2) and perfused with pre-perfusion buffer containing 0.5% sodium nitrite, 0.9% sodium chloride, and 0.1% heparin which was followed by 4% paraformaldehyde (PFA) in 0.1 M phosphate buffer. Brains were post-fixed in 4% PFA for 6-24 hours followed by 30% sucrose for 1-3 days and sliced into 40 μm sections. Tissue was permeabilized using 0.2% Triton-X 100 in 1% BSA in PBS and blocked with 1% BSA in PBS for 1 hr. Primary antibody staining was performed overnight at 4&C using rabbit anti-mouse Iba-1 (1:500; Wako Chemical Cat #019-19741) and CD68 (1:200; Bio-Rad Cat #MCA1957) with secondary staining performed for 45 minutes RT using Alexa Flour 594 donkey anti-rabbit (1:400; Invitrogen Cat #A32754, A21207), goat anti-rabbit (1:400; Invitrogen Cat #A11037), donkey anti-rat (1:400; Invitrogen A21209); Alexa Fluor 488 goat anti-rat (1:400; Invitrogen A11006), donkey anti-rabbit (1:400; Invitrogen A21206), or donkey anti-rat (1:400; Invitrogen A21208); and Alexa Flour 647 goat anti-rat (1:400; Invitrogen Cat #A21247) and mounted with DAPI Fluoromount-G medium (Southern Biotech Cat #0100-20).

Neuronal DAB staining used to quantify acute neuronal loss was performed using anti-NeuN (1:200; EMD Millipore Cat #MAB377) with secondary antibody biotin horse anti-mouse IgG (1:200; Vector Labs Cat# BA-2000). Sections were washed (0.1M PBS and 0.1M PB) and incubated with Vectastain Elite ABC-HRP Kit (1:200; Vector Laboratories PK-6100). Sections were developed for 5 min in DAB solution (3,3’diaminobenzidine; 5 mg/ml DAB, 0.1M PB, 0.000036% H_2_O_2_; Sigma-Aldrich D-5637). Acute neuronal loss in the CA1 region of the hippocampus was quantified as described previously^2^.

Neuronal Golgi staining used to visualize neurons for Sholl dendrite analyses was performed using FD Rapid Golgi Stain kit (FD Neuro Technologies), as previously described^8^. Briefly, brains were extracted and immersed in impregnation solution for 2 weeks, sectioned at 100 μm, and stained with silver nitrate solution for 10 min.

### Microscopy

The tissues used for immunohistochemistry were imaged using Zeiss LSM 880 confocal microscope (Airyscan, O.8 NA) and processed using Zen Black software (version 2.3 Sp1) and Zen Blue software (version 2.6). Microglial activation was scored as previously described^10,16^, in which microglial morphology was scored 0: <6 thin processes; 0.5: >6 but <12 thin processes; 1: 5-15 thick processes with branches, 2: 1-5 thick processes without branches; or 3: no clear processes. Microglial CD68 was scored 0: little or no expression; 1: punctate expression; 2: aggregate or punctate expression all over the cell. Composite activation scores were generated using the sum of both scores for each microglia.

Tissues used for neuron quantification and Sholl analyses were imaged using an AxioImager ZI (Zeiss, 100x oil, z=0.2-0.46 μm for neuron quantification, 40x, z=2.0 μm for dendrite measurement, 100x, z=0.5 μm for spine density) using Zen Blue software (version 3.1) and were processed using Zen Blue software (versions 2.6, 3.1). Image stacks for neuronal quantification were analyzed using StereoInvestigator programs in Neurolucida360 (MBF Bioscience). Images for Sholl analyses were analyzed on Neurolucida360 (MBF Biosciences). All raw measurements were compiled for cumulative probability distributions and analyzed by linear mixed model statistics. Dendrite lengths were collected within radii shells within neurons of mice. Dendrite length within each radial shell per neuron per mouse was the unit of observation. Radial shells were nested within neurons, which were nested within mice. A three-level linear mixed model was performed to determine whether treatment and condition were associated with dendrite length. Random intercepts for mouse and neuron were included to account of the correlation with these clusters. An interaction between treatment and condition was included to evaluate potential differential effects due to treatment and condition groups. If significant (p<0.05), pairwise comparisons were evaluated, and Bonferroni correction was applied to correct for multiple testing. If the interaction was not significant, the model was performed again, including treatment and condition as main effects. Estimated mean dendrite length by treatment and condition and corresponding 95% CIs were reported. The intra-class correlation coefficient (icc), a measure of the proportion of variation explained by neuron and mouse cluster, was also reported. The *nlme* package was used to perform the mixed model within the R program version 3.6.1 with R studio.

### Single-cell RNA-seq

Hippocampal microglia were extracted and isolated from mice as described above. Library preparation was performed according to the manufacturer’s instructions (Next GEM Single Cell 3’GEM v3.1 protocol, 10x Genomics). Briefly, microglia were resuspended in the master mix and loaded together with partitioning oil and gel beads into the chip to generate the gel bead-in-emulsion (GEM). The RNA from the cell lysate contained in every single GEM was retrotranscribed to cDNA, which contains an Illumina R1 primer sequence, Unique Molecular Identifier (UMI), and a 10x bead barcode. The pooled barcoded cDNA was then cleaned up with Silane DynaBeads, amplified by PCR, and the appropriately sized fragments were selected with SPRIselect reagent for subsequent library construction. During the library construction Illumina R2 primer sequence, paired-end constructs with P5 and P7 sequences and a sample index were added. Samples were sequenced by Genewiz (Azenta Life Sciences), 2x 150 bases on an Illumina HiSeq X 10.

FASTQ data from 10x Chromium were processed with CellRanger v6.1.2^38^ aligning to the C57BL/6 reference genome. The resulting count matrices were down sampled using the downsampleReads function in DropletUtils v1.14.2^39^, so that each mouse had the same number of reads per cell on average (based on the number of reads per cell returned by CellRanger). Empty droplets were identified with the emptyDropsCellRanger function in DropletUtils, setting the number of expected cells set to match the number estimated by CellRanger and otherwise using default parameters. The resulting matrices were combined into one joint matrix. This matrix was loaded into Seurat v4.0.0^40^ with min.features=300. The data were normalized to log counts per million (CPM) and the data were scaled, regressing out the number of genes per cell (nFeature_RNA column in the meta data). Principal Component Analysis (PCA) was calculated with RunPCA, followed by a UMAP being calculated with RunUMAP with dims=1:15 and otherwise default parameters. The data were clustered with the FindNeighbors and FindClusters functions with dims.use=1:15 and otherwise default parameters. Doublet scores were calculated with scds v1.6.0^41^. Cell types were identified using known markers (including *Snap25* for neurons, *Pdgfra* for oligodendrocyte precursor cells, *Plp1*/*Mobp* for oligodendrocytes, *Csf1r* for microglia, *Flt1* for endothelial cells, *Slc1a3*/*Gfap*/*S100b* for astrocytes, *Gad1*/*Gad2* for inhibitory neurons, and *Slc17a6*/*Slc17a7* for excitatory neurons) and using Azimuth v0.3.2^40^ with data from Allen Brain Atlas used as a reference (downloaded from https://portal.brain-map.org/atlases-and-data/rnaseq/mouse-whole-cortex-and-hippocampus-10x). Clusters consisting of non-microglia cell types and doublets were removed. This resulted in 18569 microglia split over 3 mice per condition, with 2515 Ms4a7+ microglia and 15285 Homeostatic microglia.

Cell type composition analysis was performed with the propeller.anova function in the speckle v0.0.2 package^42^, using the asin normalization and otherwise default parameters and performing Benjamini-Hochberg multiple hypothesis correction. This was followed by post hoc pairwise analyses with propeller.ttest (two-sided test based on a moderated t test) and Holm-Sidak multiple hypothesis correction. Differential expression between conditions was performed using a pseudobulk based procedure^43^, by summing up the per gene UMI counts for all cells in each sample and fitting a model with EdgeR v3.32.1^44^ using the likelihood ratio test (assumes a negative binomial model), followed by correcting p-values with fdrtool v1.2.16^45^ and performing Benjamini-Hochberg multiple hypothesis correction, and the results were reported for each comparison of interest.

Enrichment analysis was performed using fgsea v1.16.0^46^, and gene set scoring was performed with the AddModuleScores function in Seurat. Differential expression between clusters was performed using presto v1^47^ (Wilcoxon rank sum test, two-sided). The DAM and homeostatic gene sets were extracted from table S2 in Keren-Shaul et al.^26^, the NPSLE gene set was extracted from Figure 5 in Makinde et al.^29^, and the MGnD gene set was extracted from the ‘Common affected genes’ tab in Table S1 from Krasemann et al.^30^.

**Figure 5.**
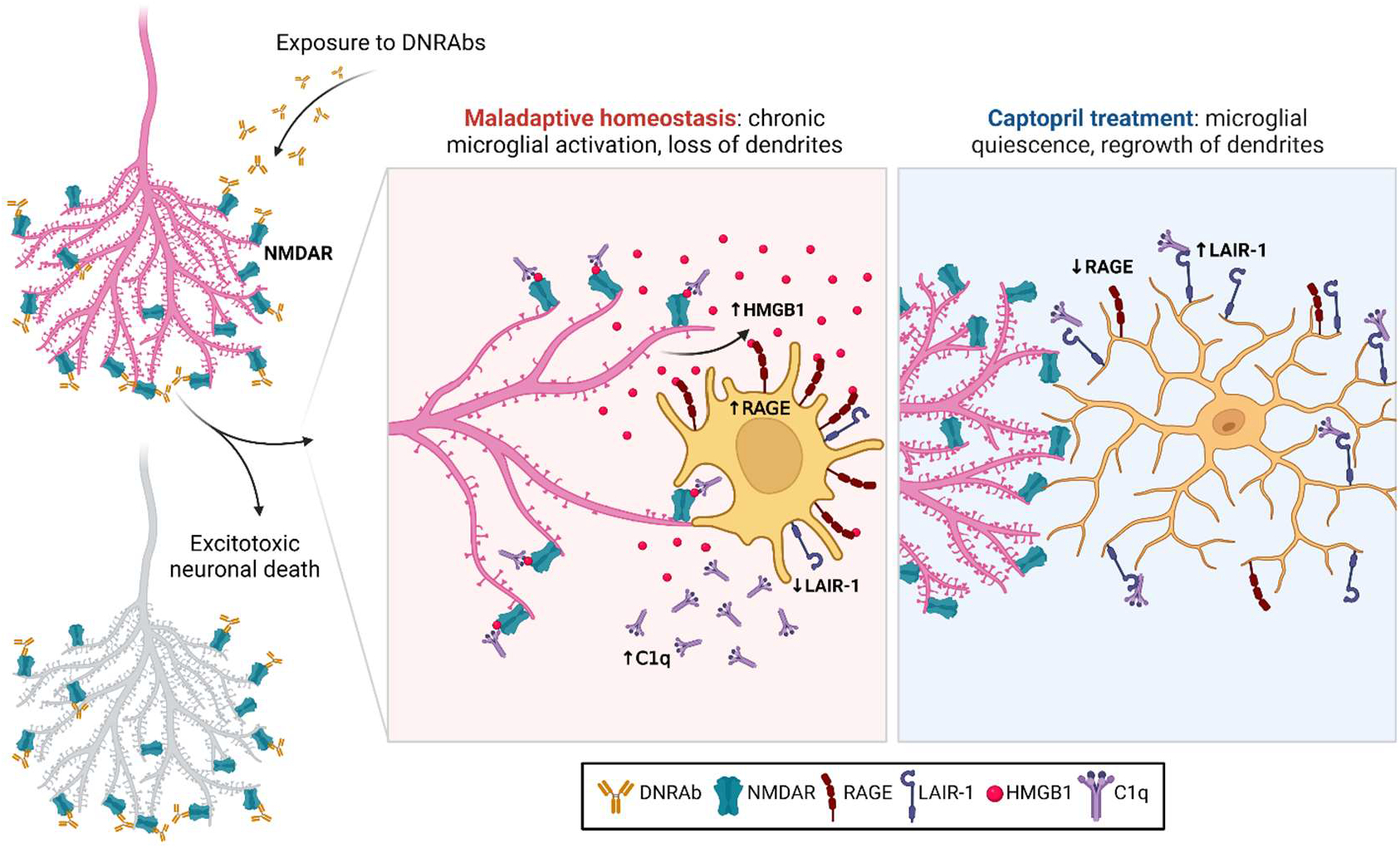
Model showing outcomes of DNRAb-mediated neuronal damage. Exposure of hippocampal CA1 pyramidal neurons to DNRAbs results in DNRAb binding to NMDARs, mediating excitotoxic death in 30% of neurons. A maladaptive equilibrium begins as a microglial response to apoptotic neuronal debris and progresses as stressed neurons secrete HMGB1, which activate microglia by binding RAGE. Activated microglia secrete proinflammatory cytokines, Type I IFN, and C1q. The secreted HMGB1 acts as a bridge by binding both NMDARs and C1q, which opsonizes synapses for microglial pruning, resulting in a loss of neuronal dendrite branching and spine density. Captopril treatment mediates microglial upregulation of LAIR-1, which induces quiescence when bound by C1q. This allows for a return to a healthy homeostasis and regrowth of dendritic branches and spines. Image created with Biorender.com.

### Statistical analyses

GraphPad Prism (version 9, GraphPad Software) was used for all statistical comparisons unless otherwise noted. ANOVA, Student’s t test, Mann-Whitney, and Kruskal-Wallis tests were performed as indicated. R (version 4.2.2) was used to perform linear mixed model analyses for Sholl analyses. All tests were performed with two-sided analysis. P>0.05 was considered statistically significant.

### Data Availability

The accession number for the sequencing data reported in this study is GEO: GSE230077 and can also be accessed through single cell portal SCP2193. Custom code can be accessed at https://github.com/seanken/LupusModel.

## Acknowledgements

We would like to thank RoseAnn Berlin and Houman Khalili for their technical assistance. This study was supported by grants from the National Institutes of Health (NIH 1P01AI073693).

## Author Contributions

K.R.C. wrote and edited the manuscript and designed, performed, and analyzed experiments. M.M. wrote and edited the manuscript and designed, performed, and analyzed experiments. S.S. wrote and edited the manuscript and designed, performed, and analyzed experiments. B.T. designed, performed, and analyzed experiments. J.W. performed and analyzed experiments. N.T. performed experiments. A.Z. performed and analyzed experiments. N.K. performed and analyzed experiments. C.K. designed, performed, and analyzed experiments and edited the manuscript. J.Z.L. designed and analyzed experiments, edited the manuscript, and oversaw the studies. B.T.V. designed, performed, and analyzed experiments, edited the manuscript, and oversaw the studies. B.D. designed and analyzed experiments, wrote and edited the manuscript, and oversaw the studies.

## Competing Interest Declaration

We have no competing interests to declare.

Supplementary Material is available for this paper.

