## Supplementary Material for "Lupus autoantibodies initiate a maladaptive equilibrium sustained by HMGB1:RAGE signaling and reversed by LAIR-1:C1q signaling"

**Supplementary Figure 1. Pathology is sustained for at least 12 months. A)** Decreased dendritic complexity in 12 month old DNRAb$^+$ compared with DNRAb$^-$ mice (mean +/- SEM; n=4-5 mice per group; n=55-59 neurons analyzed per group; linear mixed model test). **B)** Decreased dendritic spine density in 12 m.o. DNRAb$^+$ compared with DNRAb$^-$ mice (median (solid line) with quartiles (dash); n=4 mice per group; n=15-18 neurons analyzed per group; Mann-Whitney test). **C)** Representative sections of microglia in CA1 stratum radiatum stained for Iba1 (red) and CD68 (white) in 12 m.o. DNRAb$^+$ and DNRAb$^-$ B6.H2$^d$ mice (n=3 mice per group). **D)** Increased activation score in 12 m.o. DNRAb$^+$ microglia compared to 12 m.o. DNRAb$^-$ counterparts based on morphology and CD68 expression (median (solid line) with quartiles (dash); n=3 mice per group; n=110-169 microglia scored per group; Mann-Whitney test).

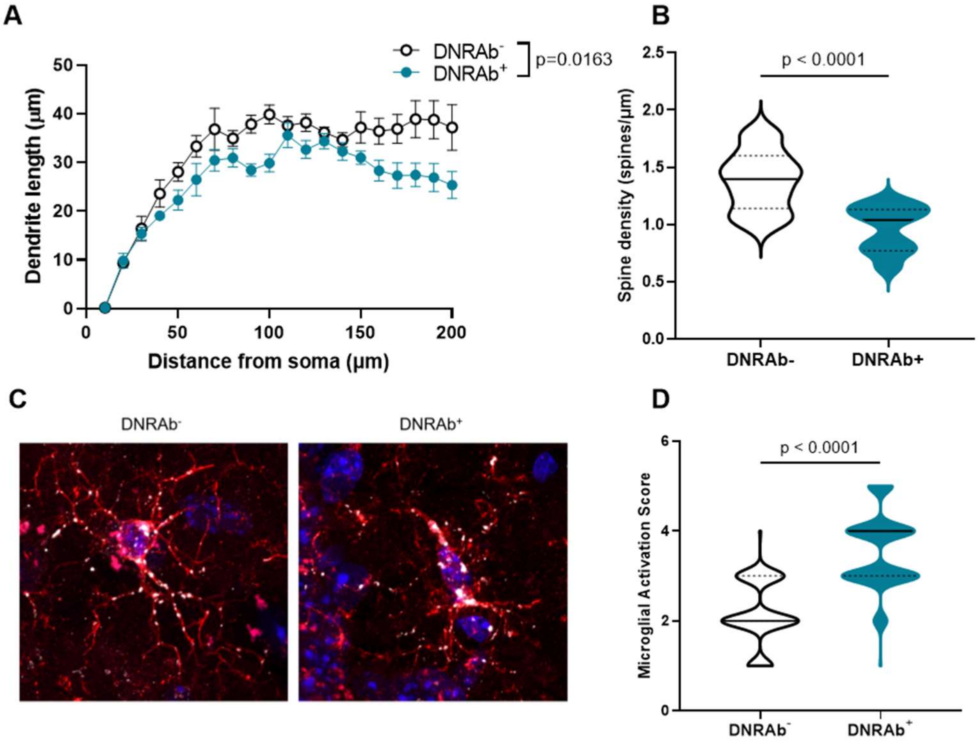

**Supplementary Figure 2. Acute neuronal loss is not affected by loss of RAGE or microglial LAIR-1. A)** Decreased CA1 neurons in WT (B6) and RAGE KO mDNRAb$^+$ mice compared to their mDNRAb$^-$ counterparts (median (solid line) with quartiles (dash); n=3-4 mice per group; n= 72-97 sections per group; Kruskal-Wallis test). B) Decreased CA1 neurons in LAIR-1 cKO DNRAb$^+$ mice compared to DNRAb$^-$ (median (solid line) with quartiles (dash); n=3 mice per group; n= 67-98 sections per group; Mann-Whitney test).

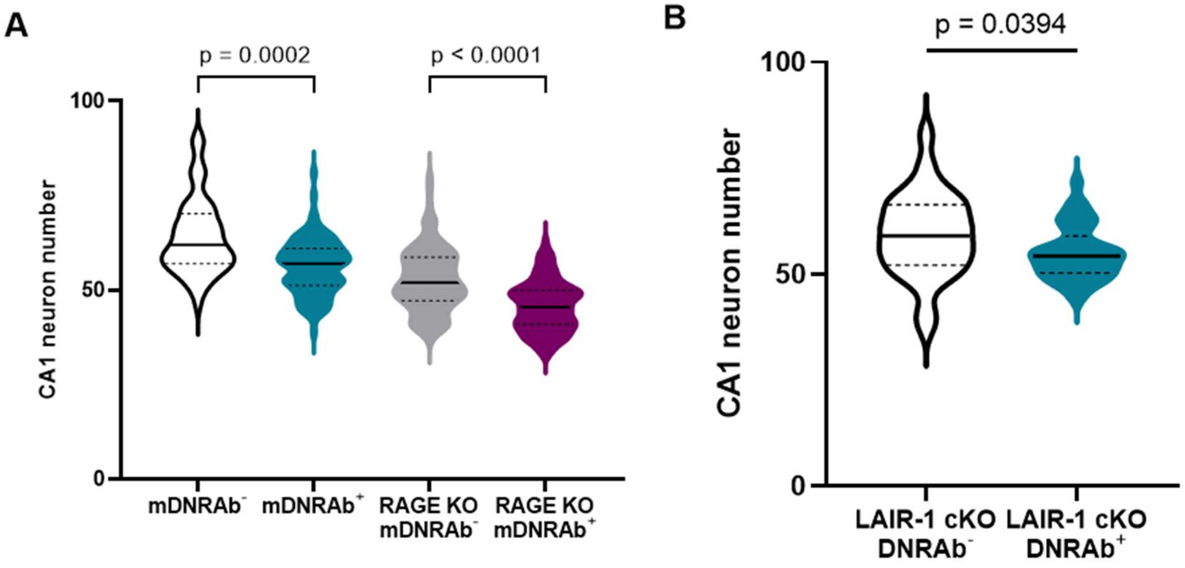

**Supplementary Figure 3. Single-cell RNA-seq clustering and quality control. A)** UMAP plot colored by clustering. **B)** UMAP plot split by mouse of origin, colored by cell type. No evidence of strong batch effects in the UMAP space. **C)** Violin plot of various QC metrics of interest, with similar distributions observed in each mouse. QC metrics include the score returned by Azimuth (Azimuth Score), number of genes per cell (nGene), number of UMI per cell (nUMI), percent UMI coming from mitochondrial reads (Percent Mitochondrial), percent UMI mapping to ribosomal proteins (Percent Ribosomal Protein), and doublet scores (scds). **D)** Feature plots of genes associated with a known microglia activation signature[48]. Subclustering within microglia subtypes is largely driven by these variables. **E)** Feature plots of number of genes per cell (nGene) and number of UMIs per cell (nUMI). Subclustering within microglia subtypes is largely driven by these variables.

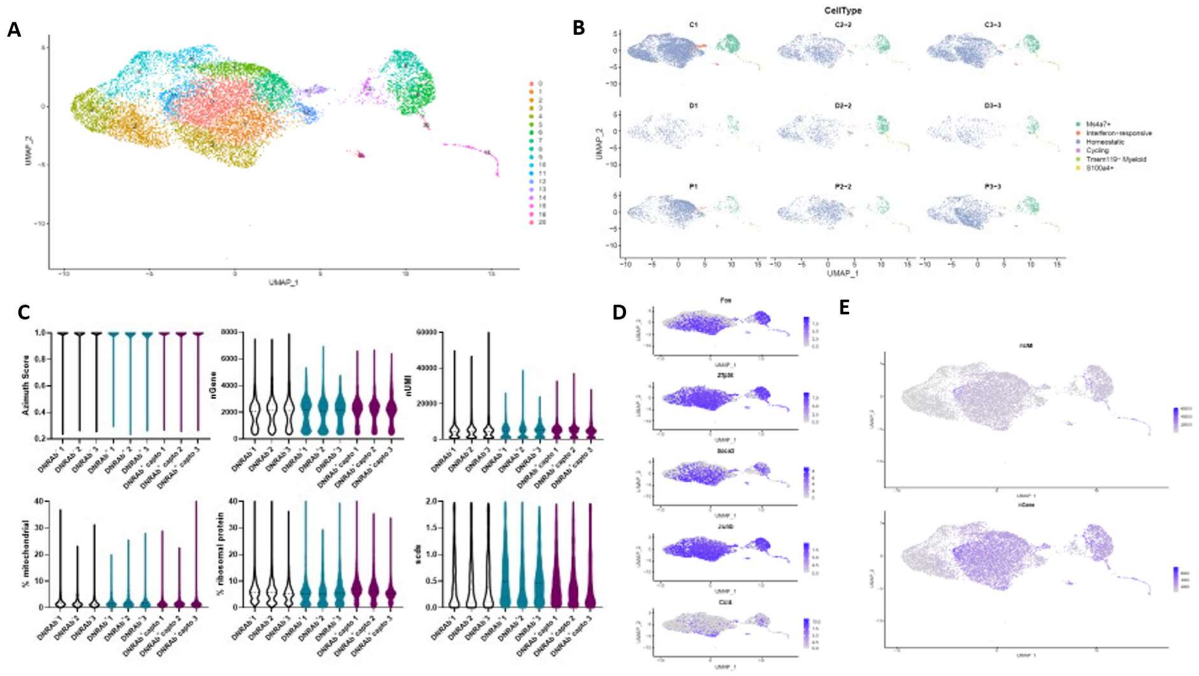

**Supplementary Figure 4. Concordance score of Ms4a7+ microglia with known microglial gene signatures. A)** Higher DAM signature gene set score in Ms4a7+ compared with Homeostatic microglia (Keren-Shaul et al. (2017)[26]; median (solid line) with quartiles (dash); n=3 mice per group; n=2515-15285 cells/cluster; Mann-Whitney test). **B)** Lower Homeostatic signature gene set score in Homeostatic compared with Ms4a7+ cluster (Keren-Shaul et al. (2017)[26]; median (solid line) with quartiles (dash); n=3 mice per group; n=2515-15285 cells/cluster; Mann-Whitney test). **C)** Higher NPSLE signature gene set score in Ms4a7+ compared with Homeostatic microglia (Makinde et al. (2020)[29]; median (solid line) with quartiles (dash); n=3 mice per group; n=2515-15285 cells/cluster; Mann-Whitney test). **D)** Higher MGnD signature gene set score in Ms4a7+ compared with Homeostatic microglia (Krasemann et al. (2017)[30]; median (solid line) with quartiles (dash); n=3 mice per group; n=2515-15285 cells/cluster; Mann-Whitney test).

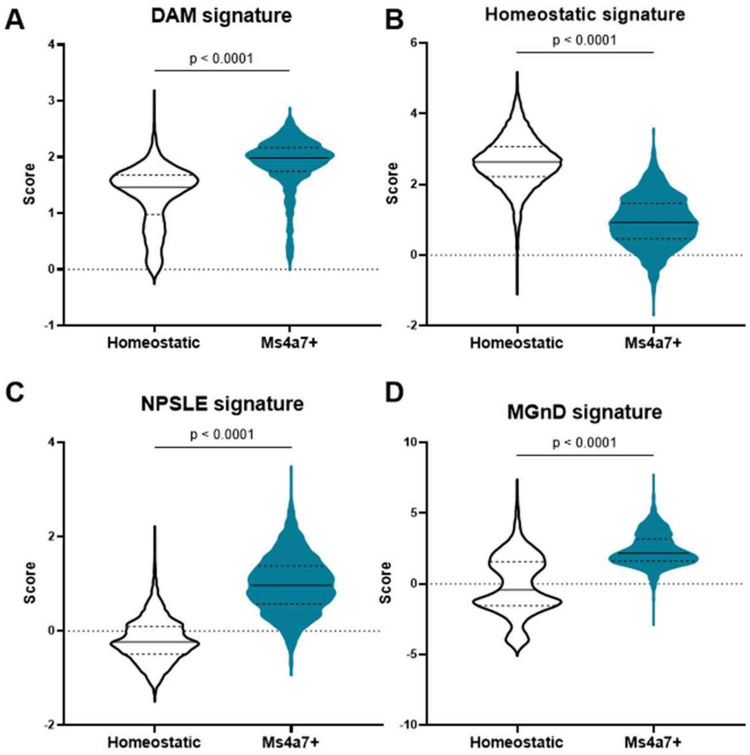

**Supplementary Table 1. Single-cell RNA-seq quality control metrics summary.** For each sample (n=3 mice per treatment group), estimated number of cells; mean reads per cell; median genes per cell; number of reads; valid barcodes; sequencing saturation; Q30 bases in barcode, RNA read, and UMI; reads mapped to genome; reads mapped confidently to genome, intergenic regions, intronic regions, exonic regions, and transcriptome; reads mapped to antisense to gene; fraction of reads in cells; total genes detected; and median UMI counts per cell.

| | $DNRAb^-$ 1 | $DNRAb^-$ 2 | $DNRAb^-$ 3 | $DNRAb^+$ 1 | $DNRAb^+$ 2 | $DNRAb^+$ 3 | $DNRAb^+$ capto 1 | $DNRAb^+$ capto 2 | $DNRAb^+$ capto 3 |
|---|---|---|---|---|---|---|---|---|---|
| Estimated Number of Cells | 6119 | 1868 | 3092 | 763 | 1257 | 754 | 2189 | 1445 | 2306 |
| Mean Reads per Cell | 24574 | 77412 | 45456 | 205512 | 130434 | 194448 | 71837 | 108980 | 66490 |
| Median Genes per Cell | 2025 | 2778 | 2504 | 2738 | 2664 | 2986 | 2815 | 2889 | 2716 |
| Number of Reads | 150370974 | 144605605 | 140548918 | 156805849 | 163955607 | 146613789 | 157251130 | 157476807 | 153326132 |
| Valid Barcodes | 0.977 | 0.976 | 0.976 | 0.975 | 0.973 | 0.974 | 0.977 | 0.975 | 0.975 |
| Sequencing Saturation | 0.48 | 0.707 | 0.595 | 0.874 | 0.8 | 0.86 | 0.73 | 0.791 | 0.696 |
| Q30 Bases in Barcode | 0.978 | 0.978 | 0.976 | 0.978 | 0.978 | 0.978 | 0.978 | 0.976 | 0.976 |
| Q30.Bases in RNA Read | 0.883 | 0.891 | 0.882 | 0.894 | 0.89 | 0.887 | 0.897 | 0.89 | 0.88 |
| Q30 Bases in UMI | 0.978 | 0.978 | 0.974 | 0.978 | 0.978 | 0.978 | 0.978 | 0.975 | 0.974 |
| Reads Mapped to Genome | 0.945 | 0.926 | 0.939 | 0.923 | 0.909 | 0.921 | 0.948 | 0.933 | 0.936 |
| Reads Mapped Confidently to Genome | 0.93 | 0.91 | 0.924 | 0.908 | 0.893 | 0.905 | 0.932 | 0.917 | 0.92 |
| Reads Mapped Confidently to Intergenic Regions | 0.042 | 0.044 | 0.043 | 0.046 | 0.047 | 0.043 | 0.043 | 0.042 | 0.04 |
| Reads Mapped Confidently to Intronic Regions | 0.365 | 0.365 | 0.378 | 0.382 | 0.379 | 0.384 | 0.387 | 0.395 | 0.392 |
| Reads Mapped Confidently to Exonic Regions | 0.522 | 0.5 | 0.504 | 0.48 | 0.467 | 0.479 | 0.503 | 0.48 | 0.488 |
| Reads Mapped Confidently to Transcriptome | 0.476 | 0.454 | 0.457 | 0.434 | 0.421 | 0.433 | 0.459 | 0.436 | 0.446 |
| Reads Mapped Antisense to Gene | 0.026 | 0.027 | 0.027 | 0.027 | 0.026 | 0.027 | 0.024 | 0.026 | 0.024 |
| Fraction Reads in Cells | 0.801 | 0.773 | 0.79 | 0.764 | 0.77 | 0.784 | 0.865 | 0.847 | 0.85 |
| Total Genes Detected | 18639 | 17196 | 17670 | 15776 | 17136 | 16388 | 17051 | 17162 | 17638 |
| Median UMI Counts per Cell | 4481 | 7153 | 5927 | 7186 | 7322 | 8300 | 7266 | 7805 | 6570 |

**Supplementary Table 2. Single-cell RNA-seq cell cluster composition frequency and percentage.** For each sample (n=3 mice per treatment group), number of cells in each cluster and the percentage of total cells this represents.

| | DNRAb⁻ 1 | DNRAb⁻ 2 | DNRAb⁻ 3 | DNRAb⁺ 1 | DNRAb⁺ 2 | DNRAb⁺ 3 | DNRAb⁺ capto 1 | DNRAb⁺ capto 2 | DNRAb⁺ capto 3 |
|---|---|---|---|---|---|---|---|---|---|
| Number of cells in Homeostatic cluster | 4924 | 1431 | 2476 | 537 | 709 | 457 | 1848 | 1119 | 1784 |
| % of total cells | 83.13355 | 80.89316 | 84.01765 | 80.63063 | 64.396 | 68.92911 | 89.57828 | 86.20955 | 83.40346 |
| Number of cells in Ms4a7+ cluster | 741 | 294 | 378 | 93 | 305 | 157 | 131 | 123 | 293 |
| % of total cells | 12.51055 | 16.61956 | 12.8266 | 13.96396 | 27.70209 | 23.68024 | 6.349976 | 9.476117 | 13.69799 |
| Number of cells in Tmem119- cluster | 41 | 17 | 38 | 15 | 50 | 18 | 23 | 33 | 34 |
| % of total cells | 0.692217 | 0.960995 | 1.289447 | 2.252252 | 4.541326 | 2.714932 | 1.114881 | 2.542373 | 1.589528 |
| Number of cells in IFN-responsive cluster | 163 | 15 | 29 | 6 | 7 | 6 | 39 | 15 | 16 |
| % of total cells | 2.751984 | 0.847937 | 0.984052 | 0.900901 | 0.635786 | 0.904977 | 1.890451 | 1.155624 | 0.748013 |
| Number of cells in Cycling cluster | 44 | 9 | 17 | 10 | 9 | 4 | 18 | 5 | 9 |
| % of total cells | 0.742867 | 0.508762 | 0.576858 | 1.501502 | 0.817439 | 0.603318 | 0.872516 | 0.385208 | 0.420757 |
| Number of cells in S100a4+ cluster | 10 | 3 | 9 | 5 | 21 | 21 | 4 | 3 | 3 |
| % of total cells | 0.168833 | 0.169587 | 0.305395 | 0.750751 | 1.907357 | 3.167421 | 0.193892 | 0.231125 | 0.140252 |

**Supplementary Table 3. Single-cell RNA-seq cell cluster comparison statistics.** For each cluster, ANOVA p-value, FDR adjusted p-value, means of the groups compared, and F statistic are reported (n=3 mice per treatment group). Posthoc pairwise t tests were performed between each treatment group within each cluster; comparisons performed, pairwise p values, and Holm-Sidak adjusted p values are reported (n=3 mice per treatment group).

| Cluster | Treatment | ANOVA p-value | FDR adjusted p-value | Mean | F statistic | Comparison | Pairwise p-value | Adjusted p-value |
|---|---|---|---|---|---|---|---|---|
| Homeostatic | DNRAb$^-$ | 0.004518 | 0.01477 | 0.82681 | 9.919631 | DNRAb$^+$ vs DNRAb$^-$ | 0.0117396 | 0.034807 |
| | DNRAb$^+$ | | | 0.71319 | | DNRAb$^+$ vs DNRAb$^+$ capto | 0.0016230 | 0.004861 |
| | DNRAb$^+$ captopril | | | 0.86397 | | DNRAb$^+$ capto vs DNRAb$^-$ | 0.2494213 | 0.577148 |
| Ms4a7+ | DNRAb$^-$ | 0.013422 | 0.026844 | 0.13986 | 6.948678 | DNRAb$^+$ vs DNRAb$^-$ | 0.0521964 | 0.148559 |
| | DNRAb$^+$ | | | 0.21782 | | DNRAb$^+$ vs DNRAb$^+$ capto | 0.0042981 | 0.012839 |
| | DNRAb$^+$ captopril | | | 0.09841 | | DNRAb$^+$ capto vs DNRAb$^-$ | 0.1673378 | 0.422693 |
| Tmem119- | DNRAb$^-$ | 0.022958 | 0.034437 | 0.00981 | 5.712187 | DNRAb$^+$ vs DNRAb$^-$ | 0.0075268 | 0.022411 |
| | DNRAb$^+$ | | | 0.03169 | | DNRAb$^+$ vs DNRAb$^+$ capto | 0.0770633 | 0.213831 |
| | DNRAb$^+$ captopril | | | 0.01749 | | DNRAb$^+$ capto vs DNRAb$^-$ | 0.1971263 | 0.482463 |
| IFN-responsive | DNRAb$^-$ | 0.488584 | 0.488584 | 0.01528 | 0.771867 | DNRAb$^+$ vs DNRAb$^-$ | 0.2552766 | 0.058696 |
| | DNRAb$^+$ | | | 0.00814 | | DNRAb$^+$ vs DNRAb$^+$ capto | 0.4146736 | 0.799463 |
| | DNRAb$^+$ captopril | | | 0.01265 | | DNRAb$^+$ capto vs DNRAb$^-$ | 0.7287750 | 0.980048 |
| Cycling | DNRAb$^-$ | 0.466644 | 0.488584 | 0.00609 | 0.82538 | DNRAb$^+$ vs DNRAb$^-$ | 0.3495265 | 0.724774 |
| | DNRAb$^+$ | | | 0.00974 | | DNRAb$^+$ vs DNRAb$^+$ capto | 0.2556850 | 0.587645 |
| | DNRAb$^+$ captopril | | | 0.00559 | | DNRAb$^+$ capto vs DNRAb$^-$ | 0.8266030 | 0.994787 |
| S100a4+ | DNRAb$^-$ | 0.004923 | 0.01477 | 0.00214 | 9.660058 | DNRAb$^+$ vs DNRAb$^-$ | 0.0039998 | 0.011952 |
| | DNRAb$^+$ | | | 0.01942 | | DNRAb$^+$ vs DNRAb$^+$ capto | 0.0033397 | 0.009985 |
| | DNRAb$^+$ captopril | | | 0.00188 | | DNRAb$^+$ capto vs DNRAb$^-$ | 0.9129096 | 0.999339 |

**Supplementary Table 4. Single-cell RNA-seq gene expression in Ms4a7+ cluster by sample.** For each sample (n=3 mice per treatment group), transcripts per million (TPM) and $log_2$TPM for each indicated gene (*Tnf, Il1b, C3ar1, Lyz2, Ifnar1, Cd93, Axl, Lair1*).

| Gene | DNRAb⁻ 1 | DNRAb⁻ 2 | DNRAb⁻ 3 | DNRAb⁺ 1 | DNRAb⁺ 2 | DNRAb⁺ 3 | DNRAb⁺ capto 1 | DNRAb⁺ capto 2 | DNRAb⁺ capto 3 |
|---|---|---|---|---|---|---|---|---|---|
| *Tnf* | | | | | | | | | |
| TPM | 78.75947 | 136.7593 | 119.0401 | 130.1744 | 139.7316 | 162.5265 | 44.29625 | 65.39778 | 154.4324 |
| $Log_2$TPM | 6.317584 | 7.106006 | 6.907372 | 7.035343 | 7.136802 | 7.353381 | 5.50132 | 6.053063 | 7.280143 |
| *Il1b* | | | | | | | | | |
| TPM | 125.6192 | 285.5573 | 69.24553 | 147.6606 | 269.6664 | 540.32 | 18.79235 | 117.716 | 233.2621 |
| $Log_2$TPM | 6.984352 | 8.16268 | 6.134335 | 7.215878 | 8.080372 | 9.080338 | 4.306871 | 6.891371 | 7.87198 |
| *C3ar1* | | | | | | | | | |
| TPM | 405.4573 | 352.4923 | 369.1798 | 462.4107 | 443.4286 | 433.7629 | 408.0625 | 417.2378 | 420.4249 |
| $Log_2$TPM | 8.66696 | 8.465535 | 8.532082 | 8.856147 | 8.795808 | 8.764085 | 8.676177 | 8.70818 | 8.719132 |
| *Lyz2* | | | | | | | | | |
| TPM | 7777.828 | 7907.481 | 7644.24 | 9634.851 | 9430.076 | 10136.92 | 6151.81 | 9920.843 | 7553.358 |
| $Log_2$TPM | 12.92534 | 12.94918 | 12.90035 | 13.2342 | 13.20321 | 13.30747 | 12.58703 | 13.27639 | 12.88309 |
| *Ifnar1* | | | | | | | | | |
| TPM | 152.459 | 178.6539 | 150.1617 | 172.9183 | 194.9023 | 213.1143 | 167.7888 | 164.8024 | 109.2552 |
| $Log_2$TPM | 7.261709 | 7.489076 | 7.239948 | 7.442266 | 7.613991 | 7.742237 | 7.399076 | 7.373321 | 6.784702 |
| *Cd93* | | | | | | | | | |
| TPM | 150.039 | 154.5765 | 175.837 | 178.747 | 199.5429 | 172.2136 | 157.0504 | 187.0376 | 85.74455 |
| $Log_2$TPM | 7.238777 | 7.281481 | 7.466276 | 7.489824 | 7.647767 | 7.436408 | 7.30424 | 7.554878 | 6.438701 |
| *Axl* | | | | | | | | | |
| TPM | 129.3591 | 152.1688 | 141.2142 | 114.6312 | 183.5588 | 220.6486 | 130.2041 | 198.8092 | 123.5459 |
| $Log_2$TPM | 7.026348 | 7.258979 | 7.151922 | 6.853387 | 7.527937 | 7.792131 | 7.035669 | 7.642479 | 6.960534 |
| *Lair1* | | | | | | | | | |
| TPM | 439.7771 | 422.7982 | 430.645 | 415.781 | 407.8512 | 389.6332 | 437.5933 | 395.0026 | 441.6305 |
| $Log_2$TPM | 8.783905 | 8.727234 | 8.753701 | 8.703146 | 8.675432 | 8.609671 | 8.77674 | 8.629366 | 8.789959 |

**Supplementary Table 5. Single-cell RNA-seq Ms4a7+ cell gene expression comparison statistics.** For each gene, comparisons between each group were performed using a likelihood ratio test with FDR and Benjamini-Hochberg corrections. LogFC, logCPM, LR, p-value, and adjusted p-value (padj) are reported (n=3 mice per treatment group).

| Gene | Comparison | logFC | logCPM | LR | p-value | padj |
|---|---|---|---|---|---|---|
| *Tnf* | DNRAb$^+$ vs DNRAb$^-$ | 0.475008 | 6.977014 | 6.651073 | 0.00991 | 0.151184 |
| | DNRAb$^+$ vs DNRAb$^+$ capto | 0.756772 | 6.935088 | 5.489881 | 0.019127 | 0.32575 |
| | DNRAb$^+$ capto vs DNRAb$^-$ | -0.29284 | 6.696582 | 0.733314 | 0.391811 | 0.993521 |
| *Il1b* | DNRAb$^+$ vs DNRAb$^-$ | 1.092901 | 7.897332 | 6.787678 | 0.009179 | 0.143578 |
| | DNRAb$^+$ vs DNRAb$^+$ capto | 1.467294 | 7.871111 | 5.907503 | 0.015077 | 0.287376 |
| | DNRAb$^+$ capto vs DNRAb$^-$ | -0.36953 | 7.197758 | 0.343762 | 0.557666 | 0.993521 |
| *C3ar1* | DNRAb$^+$ vs DNRAb$^-$ | 0.32206 | 8.688049 | 7.359287 | 0.006672 | 0.116852 |
| | DNRAb$^+$ vs DNRAb$^+$ capto | 0.154019 | 8.78866 | 0.790979 | 0.373804 | 0.892005 |
| | DNRAb$^+$ capto vs DNRAb$^-$ | 0.179949 | 8.638248 | 1.250056 | 0.263542 | 0.993521 |
| *Lyz2* | DNRAb$^+$ vs DNRAb$^-$ | 0.403402 | 13.11144 | 12.55886 | 0.000394 | 0.015454 |
| | DNRAb$^+$ vs DNRAb$^+$ capto | 0.345297 | 13.14444 | 2.232687 | 0.135119 | 0.701726 |
| | DNRAb$^+$ capto vs DNRAb$^-$ | 0.069946 | 12.95071 | 0.097732 | 0.754569 | 0.998601 |
| *Ifnar1* | DNRAb$^+$ vs DNRAb$^-$ | 0.37003 | 7.470336 | 7.467909 | 0.006281 | 0.111822 |
| | DNRAb$^+$ vs DNRAb$^+$ capto | 0.465003 | 7.461662 | 4.029725 | 0.044705 | 0.475877 |
| | DNRAb$^+$ capto vs DNRAb$^-$ | -0.09564 | 7.275785 | 0.190183 | 0.662764 | 0.993521 |
| *Cd93* | DNRAb$^+$ vs DNRAb$^-$ | 0.295406 | 7.430908 | 5.597676 | 0.017984 | 0.21385 |
| | DNRAb$^+$ vs DNRAb$^+$ capto | 0.410382 | 7.39705 | 2.13733 | 0.143752 | 0.71292 |
| | DNRAb$^+$ capto vs DNRAb$^-$ | -0.11804 | 7.253963 | 0.183124 | 0.668702 | 0.993521 |
| *Axl* | DNRAb$^+$ vs DNRAb$^-$ | 0.417811 | 7.310917 | 5.867303 | 0.015425 | 0.193616 |
| | DNRAb$^+$ vs DNRAb$^+$ capto | 0.267126 | 7.402549 | 0.919316 | 0.337655 | 0.876081 |
| | DNRAb$^+$ capto vs DNRAb$^-$ | 0.137412 | 7.193658 | 0.380254 | 0.537467 | 0.993521 |
| *Lair1* | DNRAb$^+$ vs DNRAb$^-$ | -0.01522 | 8.720331 | 0.018162 | 0.892797 | 0.979846 |
| | DNRAb$^+$ vs DNRAb$^+$ capto | -0.01919 | 8.731565 | 0.012642 | 0.910478 | 0.991134 |
| | DNRAb$^+$ capto vs DNRAb$^-$ | 0.013904 | 8.752519 | 0.008258 | 0.927592 | 0.998601 |
